# Increased adaptive potential in novel environments can be predicted from genetic variance in development time expressed in native environments

**DOI:** 10.1101/2021.02.04.429835

**Authors:** Greg M. Walter, Keyne Monro, Alastair Wilson, Delia Terranova, Enrico la Spina, Mari Majorana, Giuseppe Pepe, Sarah du Plessis, James Clark, Salvatore Cozzolino, Antonia Cristaudo, Simon J. Hiscock, Jon Bridle

## Abstract

While many organisms shift their development plastically to maintain fitness as environments change, such plasticity has limits. Population mean fitness is expected to decline when novel environments exceed the limits to plasticity, but the level of fitness costs is expected to vary among genotypes that can increase adaptive potential. We lack fundamental insights into how and when changes in early development traits increase adaptive potential in novel environments, which limits our ability to predict the response of natural populations to global change. To test whether genetic variation in development time is associated with increased adaptive potential in novel environments, we used a breeding design to generate c.20,000 seeds of two ecologically contrasting Sicilian species of daisies (*Senecio,* Asteraceae) adapted to high and low elevations on Mount Etna. We planted the seeds across four elevations that included the native range of each species, and a novel elevation. We tracked seedling mortality and measured development time as the number of days it took seedlings to establish. As predicted, genetic variance in survival increased at novel elevations. However, genetic variance in development time showed the opposite trend, decreasing at novel elevations. A strong negative genetic correlation between development time in the native range and survival at novel elevations suggested that genotypes with faster development in native environments survived better in novel environments. These results were consistent across the two ecologically contrasting species, suggesting that genetic variance in early development in native environments could be used to predict population responses to novel environments.

## Introduction

As global change creates increasingly novel environments, natural populations will need to shift their geographical ranges or adapt if they are to persist (Bridle and Hoffmann 2022; Urban et al. 2024). Plasticity, the ability of genotypes to express different phenotypes in different environments, provides an immediate response that can help populations to maintain fitness as environments change (Via 1993; Via et al. 1995; Charmantier et al. 2008). In particular, plasticity during early development can determine whether juveniles reach maturity and are able to reproduce in new environments (West-Eberhard 2003; Sultan 2007; Snell-Rood et al. 2018; Smallegange 2022). Plasticity is adaptive when it helps genotypes to buffer familiar variation in the environment, but should fail to maintain high fitness (i.e., become nonadaptive) in novel environments (Ghalambor et al. 2007). Theory predicts that where adaptive plasticity fails and population mean fitness declines, variation among genotypes should increase as they vary in their sensitivity to the new environment, which increases the adaptive potential of the population (Hermisson and Wagner 2004; Lande 2009; Chevin et al. 2010). The genotypes that suffer less (i.e., perform relatively better than other genotypes) in novel environments can help prevent more severe fitness declines for the population, which could then aid population persistence (Chevin and Bridle 2025). Understanding how genetic differences in life history and fitness emerge when the limits to adaptive plasticity are exceeded remains a fundamental challenge to predicting the adaptive capacity of natural populations facing global change.

Although plasticity in early development is widespread, it is not necessarily adaptive in novel environments (Ghalambor et al. 2007; Ghalambor et al. 2015; Acasuso-Rivero et al. 2019). Biophysical constraints create developmental limits that determine the capacity for adaptive plasticity during early development, which can then influence ecological limits (Angert et al. 2008; Angert et al. 2014; Eriksson and Rafajlovic 2022; Khare et al. 2024). For example, while increases in temperature favour faster development, physiological limits will determine the limit to how fast development can occur, and drastic changes in development rate are likely to incur trade-offs with survival, development or maintenance (Arendt 1997; Willi and Van Buskirk 2022). This means that although plasticity may allow genotypes to drastically change their developmental traits in novel environments, even to the extent that they match the development of plants native to that environment, such plasticity can incur other fitness costs that reduce survival (Van Kleunen and Fischer 2005; Walter et al. 2018; Cotto et al. 2019; Schneider 2022). Where plasticity during early development no longer maintains fitness, the extent to which genotypes vary in their developmental responses to novel environments could then determine population persistence (Nussey et al. 2005; Ghalambor et al. 2007; Hoffmann and Sgrò 2011). However, we lack experiments that connect genetic variance in fitness with early life history traits in natural environments, especially as conditions reach, and then exceed, current ecological margins (Angert et al. 2014; Angert et al. 2020).

Genetic differences in plasticity emerge as genotype-by-environment interactions (G×E), which manifest as a change in rank order of the genotypes across environments (**Fig. 1a**), and/or as a change in among-genotype variance across environments (**Fig. 1b**; Mackay 1981; Via and Lande 1985). Although we know that patterns of G×E change depending on the trait and environmental context (Merilä and Fry 1998; Sultan 2007; Culumber et al. 2015; Saltz et al. 2018), we have a poor understanding of whether G×E differs for developmental traits compared to fitness related traits, particularly under field conditions. Linking genetic variation in early life history with adaptive potential as environments become novel is therefore challenging, but crucial if we are to understand the adaptive capacity of populations and ecological communities that are being pushed beyond their current ecological margins (Anderson et al. 2012).

**Fig. 1.**
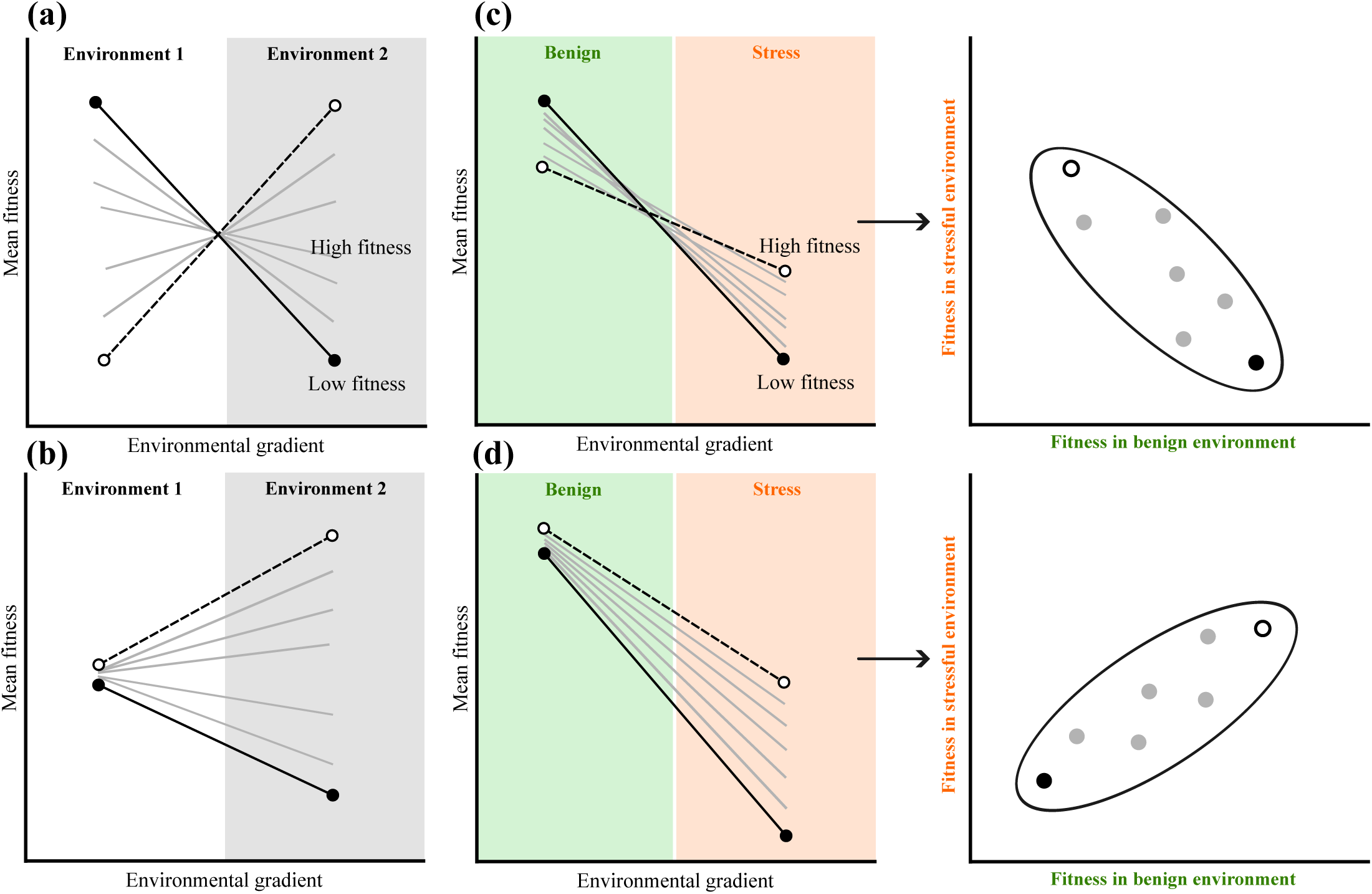
Genotype-by-environment interactions (G×E) for hypothetical genotypes that are represented by different lines, and with emphasis on the extreme differences in responses (dashed lines with open circles versus solid lines with closed circles). G×E can occur across environments as: **(a)** changes in rank of genotypes across environments, or **(b)** as changes in variance among genotypes across environments. **(c-d)** When faced with a stressful environment that reduces mean fitness, G×E means that some genotypes could suffer less under stress (dashed lines with open circles), while other genotypes suffer large reductions in fitness (solid lines with closed circles). This produces the same pattern of G×E as in **(a-b)**, but includes reductions in mean fitness. Right-hand panels in **c-d** show the particular case where a change in genotype rank produces a negative genetic correlation across environments, whereas a change in variance produces a positive correlation. Note that it is possible for a combination of **(c)** and **(d)** to determine the expression of genetic variance in fitness in novel environments.

Genetic differences in fitness and developmental sensitivity to the environment can be created by a combination of both forms of G×E. Genetic correlations in fitness between native and novel environments can weaken or even become negative as environments become novel, in the latter case genotypes with high fitness in novel environments have lower fitness in native environments (**Fig. 1c**; Angert et al. 2008; Walter et al. 2023). Novel environments themselves may also induce the expression of hidden genetic variation that increases genetic variance in fitness in novel environments (**Fig. 1d**; Hermisson and Wagner 2004; Paaby and Rockman 2014; Shaw and Shaw 2014; Walter et al. 2023). Fisher’s (1930) fundamental theorem of natural selection quantifies the potential for adaptation as the amount of genetic variance in relative fitness, 𝑉_𝐴_(𝜔), which is the ratio of genetic variance in absolute fitness to mean absolute fitness. Evidence suggests that adaptive potential changes with the environment (Sheth et al. 2018), and that natural populations harbour genetic variance in fitness within their native range (Hendry et al. 2018; Kulbaba et al. 2019; Peschel et al. 2020; Bonnet et al. 2022). However, we have a limited understanding of how 𝑉_𝐴_(𝜔) changes across environmental gradients. Although some evidence suggests that 𝑉_𝐴_(𝜔) – and therefore adaptive potential – can increase in novel or stressful environments (Torres-Martínez et al. 2019; Walter et al. 2023), meta-analyses remain equivocal (Hoffmann and Merilä 1999; Charmantier and Garant 2005; Hendry et al. 2018). We therefore lack data from field studies with multiple species to understand how genetic variance in early life history determines fitness across an ecological gradient that includes native and novel elevations.

While theory suggests that G×E should increase genetic variance in fitness in novel environments, predictions for how G×E emerges in early life history stages are more difficult. Strong selection during early development often removes unfit genotypes before they can be measured, which means that quantifying genetic variation in early life history traits is challenging (Hadfield 2008; Kruuk et al. 2008; Wadgymar et al. 2017). As a consequence, although we know that evolutionary potential in early life history traits can change with the environment (Chantepie et al. 2024), we still do not know whether genetic variance in early life history traits increases in novel environments (Charmantier and Garant 2005; Garant et al. 2008; Wood and Brodie III 2015; Riley et al. 2023). Naively, we would predict that because genotype-specific changes in early development should have consequences for fitness, we would expect increased genetic variance in fitness in novel environments to be correlated with a similar increase in genetic variance in early development traits. To predict the response of natural populations facing global change, we therefore need to understand how G×E in early life history traits changes across environments, and whether such changes determine adaptive potential in novel environments (Chirgwin et al. 2021).

To test whether genetic variance in development time and fitness increases where adaptive plasticity fails to maintain fitness in novel environments, we generated seeds of two closely related species of Sicilian daisy (*Senecio*, Asteraceae), which we planted along an elevational gradient on Mt. Etna. *Senecio chrysanthemifolius* is a short-lived perennial with dissected (more complex) leaves that occupies disturbed habitats (e.g. roadsides) at 400-1500m.a.s.l (metres above sea level) on Mt. Etna. By contrast, *Senecio aethnensis* is a longer-lived perennial with entire glaucous (simpler and waxy) leaves endemic to lava flows above 2000m.a.s.l on Mt. Etna, where individuals grow back each spring after being covered by snow in winter (Walter et al. 2020). Species also differ in plasticity in leaf traits, with *S. chrysanthemifolius* exhibiting a greater capacity for adaptive plasticity when faced with novel environments compared to *S. aethnensis* (Walter et al. 2022; Walter et al. 2024). Here, we use data derived from a seed transplant experiment that tested how genetic variance in multivariate leaf phenotypes changed across elevations (Walter et al. 2024). In this previous study, we found that changes in genetic variance were associated with low adaptive potential for the high-elevation *S. aethnensis* seedlings planted at low elevations. This contrasted with the low-elevation *S. chrysanthemifolius* seedlings, which displayed smaller changes in genetic variance when planted at higher elevations, and greater potential to adapt to the novel high elevation.

We used a quantitative genetic breeding design to produce seeds for c.100 families of each species, which we reciprocally planted across an elevational gradient that included the native elevation of each species and two intermediate elevations. For seedlings that emerged, we tracked mortality, and measured development time as the time it took seedlings to establish (produce 10 leaves). With these data, we first tested for patterns of adaptive divergence in survival, emergence and seedling establishment between the native habitats. We then tested three hypotheses to understand how genotypes of each species varied in development time and survival across the ecological gradient. *<u>Hypothesis I:</u>* If development time is important for maintaining survival across elevations, we expected selection on development time to change across elevations similarly for both species. *<u>Hypothesis II:</u>* As environments become novel and plasticity can no longer maintain fitness, we predicted that declines in survival would be associated with stronger patterns of G×E in both development time and survival. We therefore expected that as environments became novel, genetic variance would increase, and genetic correlations would change from strongly positive (+1) between environments within the native range, to weakly positive (e.g., 0.2-0.5) or negative between native and novel environments. *<u>Hypothesis III:</u>* If patterns of G×E in development time are linked to survival as environments become novel, we would detect strong genetic correlations between development time and survival, but only for species at novel elevations.

## Methods and materials

### Breeding design and field experiment

To produce seeds of both species, we collected cuttings from c.80 mature individuals from natural populations of both species, and we propagated one cutting from each individual in the glasshouse. When plants produced buds, we covered branches with perforated bread bags to exclude pollinators while providing adequate airflow. For each species, we randomly assigned half of the individuals (which are hermaphroditic and self-incompatible) as females (dams), the other half as males (sires). We then used a factorial design to randomly mate sires to dams in 3×3 blocks (three sires mated to three dams per block; **Fig. 2a**) to produce 94 families of *S. aethnensis* (*n*=36 sires, *n=*35 dams) and 108 families of *S. chrysanthemifolius* (*n=*38 sires, *n*=38 dams). We performed crosses by removing flowerheads from sires and rubbing their anthers on flowerheads of dams. Fertilised flowerheads were labelled and covered with mesh bags to catch seeds once mature. Due to limitations in glasshouse space, we performed crosses on species separately. For *S. chrysanthemifolius*, we sampled individuals from five independent locations at the base of Mt. Etna in June-July 2017 (**Table S1** and **Fig. S1**), and between January and February 2018. For *S. aethnensis*, we sampled individuals from a range of elevations on both the North and South of Mt. Etna in October 2018 (**Table S1** and **Fig. S1**), and conducted crosses between January and March 2019.

**Fig. 2.**
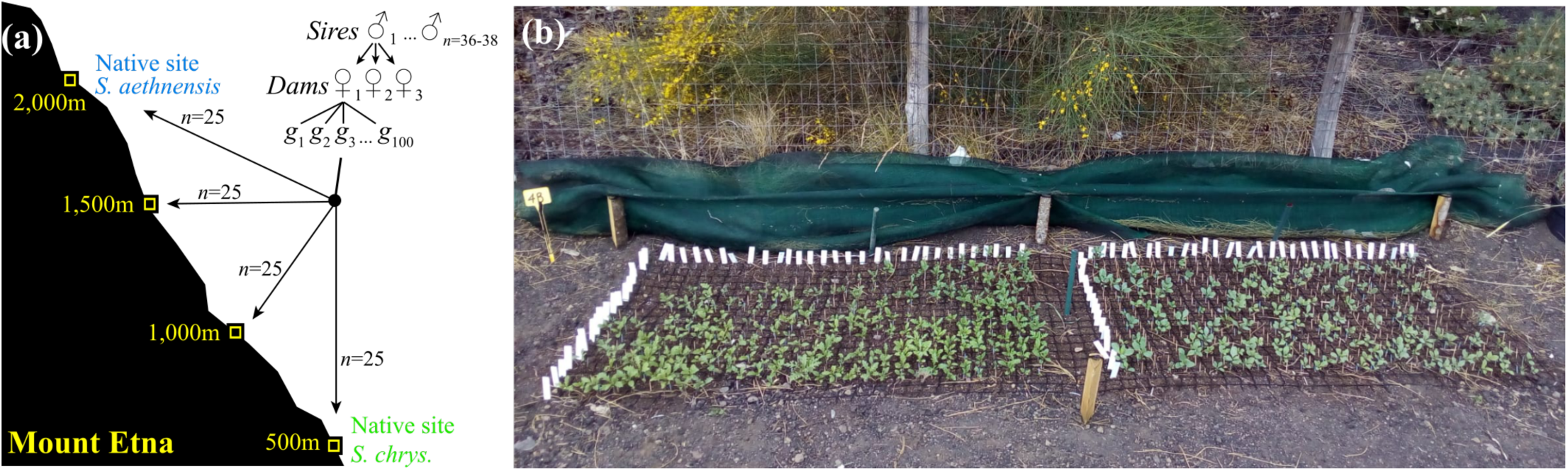
The experimental design. **(a)** For two Etnean *Senecio* species, we mated 36-38 sires each to 3 dams, and produced 100 seeds from each mating (family). We planted 25 seeds from each family at each of four elevations that represent the native site of each species and two intermediate elevations, and tracked the emergence of each seedling, development time to establishment, and survival. **(b)** Photo of an experimental block with 8-week-old seedlings of both species at 2,000m (*S. chrysanthemifolius* on left).

In April 2019, we planted 25 seeds per family at each of four elevations on Mt. Etna that included the native habitats of both species (500m and 2,000m) and two intermediate elevations (1,000m and 1,500m) (**Fig. 2b**; Walter et al. 2022; Walter et al. 2024). Vagrant individuals of these species are often found between 1,000-1,500m, which means these intermediate elevations could support range expansions. At each elevation, we randomised seeds of each family into five experimental blocks (*S. aethnensis n*=432 seeds/block, *n=*2,160 seeds/site; *S. chrysanthemifolius n=*540 seeds/block, *n=*2,700 seeds/site; Total N=19,232 seeds). However, *S. aethnensis* grew slowly and produced too few flowers in the glasshouse to obtain 100 seeds for each family, meaning that four, eight and ten families were represented by 15, 20 and 25 seeds per elevation, respectively.

To prepare each experimental block, we cleared the ground of plant matter and debris, and placed a plastic grid (4cm square cells) on the ground. We attached each individual seed to the middle of a toothpick using non-drip super glue and then pushed each toothpick into the soil of a single grid cell so that the seed sat 1-2mm below the soil surface (Walter et al. 2016). To replicate natural germination conditions, we suspended 90% shade-cloth 20cm above each plot and kept the seeds moist until seedling emergence ceased (2-3 weeks). After this initial period, we replaced the shade-cloth with a lighter 40% shade-cloth to replicate shade that naturally-growing plants are often found under. This maintained a strong environmental effect at each elevation without exposing plants to extreme environmental conditions. For the seedlings that emerged, we focused on recording mortality and the time it took seedlings to establish (produce ten leaves). The experiment concluded in January 2020 when mortality stabilised at all sites (**Fig. S2**) and plants started growing into each other, which increased competition and would have likely biased further data collection.

### Statistical analyses

All analyses below were performed in R version 3.6.1 (R Core Team 2024). Throughout the analyses, we treated elevation as categorical (rather than continuous) so that we could test how development time and survival change across elevation as environments move from native to novel. First, we assess how the *Senecio* species differed in emergence, survival and development across the elevation gradient, and we then test the three hypotheses described in the introduction.

### Quantifying mean differences in seedling traits across elevations

To test how both species emerged, survived and developed across the elevation gradient, we applied the linear mixed model

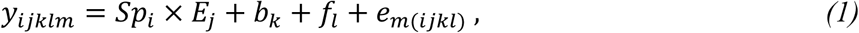

where the fixed effects include 𝑆𝑝_𝑖_ × 𝐸_j_as the *i*th species (*Sp*) planted at the *j*th elevation (*E*) included as categorical variables. Random effects include 𝑏_𝑘_ as the *k*th experimental block, 𝑓_𝑙_ as the *l*th family within species, and 𝑒_𝑙(𝑖*j*𝑘)_represents the residual. We used type III ANOVA to test for significant elevation×species interactions, and *emmeans* (Lenth 2019) to obtain marginal means for each species at each elevation.

We applied equation 1 separately for emergence, seedling establishment and survival to the end of summer, with each variable included as a binary response variable (𝑦_𝑖j𝑘𝑙_) using a logit link function with *glmmTMB* (Brooks et al. 2017). For each trait, equation 1 estimated the species mean at each elevation. Seedling establishment and survival excluded seeds that failed to germinate and emerge, and so only represent fitness of the seedlings that successfully emerged. However, preliminary analyses suggested that including variation in emergence success did not change our results or their interpretation. To understand how development time to seedling establishment changed across elevations, we applied equation 1 with development time as a gaussian-distributed response variable.

### Hypothesis I: Selection on development time changes across elevations

To quantify selection on development time for each species and elevation, we used *glmmTMB* to apply the generalised linear mixed model

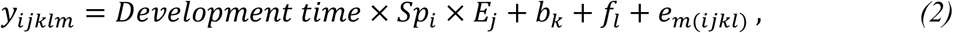

where fixed effects include the interaction between development time, species and elevation. We included survival as the binomially-distributed response variable (𝑦_𝑖j𝑘𝑙𝑚_) with a probit link function. For the seedlings that successfully established, this estimates directional selection on development time based on differences in survival. Random effects include 𝑏_𝑘_ as the *k*th block and 𝑓_𝑙_ as the *l*th family within species, and 𝑒_𝑚(𝑖*j*𝑘𝑙)_ is the residual. We then transformed the coefficients from the link to the data scale and divided them by the mean survival so that they represent selection gradients (Janzen and Stern 1998). A significant three-way interaction (𝐷𝑒𝑣𝑒𝑙𝑜𝑝𝑚𝑒𝑛𝑡 𝑡𝑖𝑚𝑒 × 𝐸𝑙𝑒𝑣𝑎𝑡𝑖𝑜𝑛 × 𝑆𝑝_𝑖_) would suggest that selection on development time differs between species and changes across elevation.

### Hypothesis II: G×E in survival and development time increases in novel environments

To estimate additive genetic variance in both survival and development time, we used *MCMCglmm* (Hadfield 2010) to apply

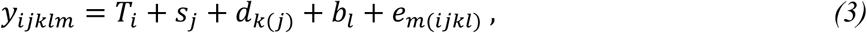

where transplant elevation (𝑇_𝑖_) was the only fixed effect. Random effects included 𝑠_j_ as the *j*th sire, 𝑑_𝑘(j)_ the *k*th dam nested within sire, 𝑏_𝑙_ the variance among blocks within a transplant elevation, and 𝑒_𝑚(𝑖*j*𝑘𝑙)_ the residual variation. We use a nested approach to estimate additive genetic variance. This reduces the number of parameters estimated by omitting the sire×dam interaction, which becomes incorporated into the residual variance without inflating the additive genetic variance (Schielzeth and Nakagawa 2013; Walter et al. 2024). We then quantified additive genetic variance as four times the sire variance (Lynch and Walsh 1998). By including transplant elevation as a fixed effect and specifying unstructured covariance matrices for the sire component, we estimated a 4×4 matrix with variance at each elevation along the diagonal and covariances among elevations on the off-diagonals. To estimate the genetic correlations between elevations, we transformed the covariances into correlations using the *cov2cor* function.

We used equation 3 to estimate genetic variance in survival and development time separately. Response variables (𝑦_𝑖j𝑘𝑙𝑚_) therefore included development time (gaussian) and survival (binary) as the presence/absence of plants after mortality stabilised at the end of summer (28^th^ of August; **Fig. S2**). For survival we specified a binomial error distribution with a probit link function. We used chains with a burn-in of 150,000 iterations, a thinning interval of 1,500 iterations and saving 1,000 iterations that provided the posterior distribution for all parameters estimated. We confirmed model convergence by checking the chains mixed sufficiently well, that autocorrelation was lower than 0.05, and that our parameter-expanded prior was uninformative (Hadfield 2010). Given that *MCMCglmm* constrains variances to be positive, we tested whether our estimates of genetic variance were statistically significant by comparing our observed estimates to a null distribution, which we created by randomly shuffling the survival and development time data among the families 200 times and re-estimating genetic variance for each randomised dataset (Walter et al. 2024). Where our observed estimates exceeded the null distribution provides evidence that we captured statistically significant genetic variation for a given trait estimated in a given environment.

To quantify changes in genetic variance in survival across elevations, we first back-transformed estimates of variance from the link scale to the data scale using *QGlmm* (de Villemereuil et al. 2016). For both traits, we then calculated genetic variance in relative survival and development time at each elevation by dividing absolute variance for each trait by the squared mean estimate of each trait. This represents genetic variance relative to the mean at each transplant elevation, and for survival this represents adaptive potential (Houle 1992; Hansen et al. 2011).

### Hypothesis III: G×E in development time is associated with survival in novel environments

To quantify genetic correlations between development time and survival, we used *MCMCglmm* to apply equation 3, but using the data collected within transplant elevation (and so removed the fixed effect of transplant elevation). Instead, we applied a bivariate model that included both development time and survival as response variables. By specifying an unstructured covariance matrix for the sire component, we estimated (for each species) a 2×2 matrix representing genetic variance in each trait and the genetic covariance between the traits. We then converted the covariances to correlations to quantify the strength of genetic correlations between development time and survival.

## Results

### Evidence of adaptive divergence between species from elevational extremes

Across all elevations, average seedling emergence was 75-85% for *S. chrysanthemifolius* and 50-70% for *S. aethnensis* (**Fig. 3a**). For *S. aethnensis,* c.60% of seedlings established at all elevations, which contrasted with *S. chrysanthemifolius*, where 75% of seedlings established at the home site but gradually reduced across elevations to 25% at the 2,000m elevation (**Fig. 3b**). Mortality stabilised after summer (**Fig. S2**), at which point evidence of adaptive divergence between the two species emerged. Both species showed greater probability of survival within their native range compared to a novel elevation, which is the native habitat of the other species (species×elevation *χ^2^*(3)=978.71, P<0.001; **Fig. 3c**). At each of the native habitats (500m and 2000m), the native species survived better than the species from the other habitat (**Fig. 3c**). Therefore, *S. chrysanthemifolius* survived better than *S. aethnensis* at low elevations (500m Z=18.7, P<0.001; 1,000m Z=3.2, P=0.0014), and *S. aethnensis* survived significantly better at higher elevations (1,500m Z=8.4, P<0.001; 2,000m Z=14.9, P<0.001).

**Fig. 3.**
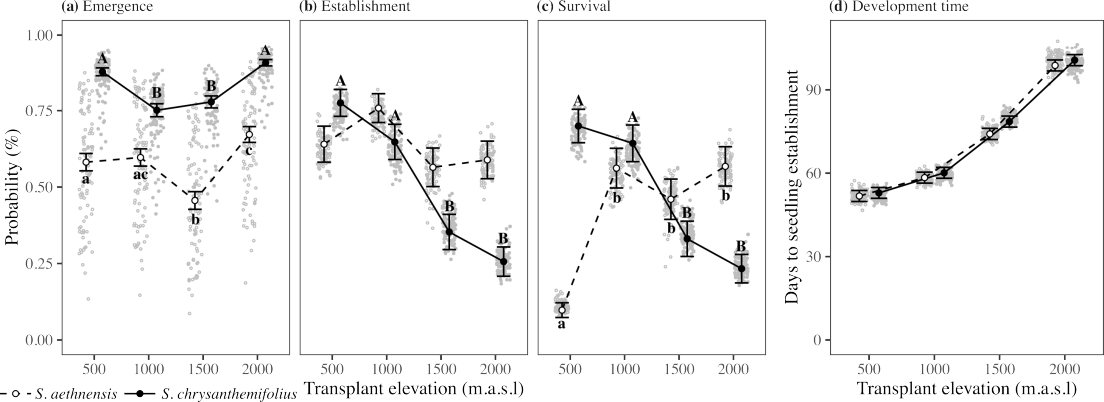
Changes in seedling traits for the two Etnean *Senecio* species planted across an elevation gradient. Lowercase and uppercase letters denote significant differences (α=0.05 with Bonferroni adjustment) across elevations for *S. aethnensis* (open circles and dashed lines) and *S. chrysanthemifolius* (closed circles and solid lines), respectively. **(a)** Seedling emergence was greater for *S. chrysanthemifolius,* which was greater at the elevational extremes. Emergence was greatest for *S. aethnensis* at its native elevation (2,000m). **(b)** The probability of seedling establishment remained similar across elevations for *S. aethnensis*, but was reduced at higher elevations for *S. chrysanthemifolius.* **(c)** As evidence of adaptive divergence, both species survived to the end of summer (end of September) better at their home site than the foreign species, and better at their home site than at the novel elevation. **(d)** Development time was slower at higher elevations (all elevations were significantly different from each other) with *S. aethnensis* showing slightly faster development than *S. chrysanthemifolius*.

### Hypothesis I: Selection on development time changes across elevations similarly for both species

Development time changed significantly across elevations (*χ^2^*(3)=357.39, P<0.001), with seedlings establishing faster at lower elevations (**Fig. 3d**). A significant interaction between development time and elevation suggested that species responded differently to elevation (*χ^2^*(3)=9.06, P=0.0285). Seedlings of *S. aethnensis* tended to establish earlier than *S. chrysanthemifolius,* particularly at higher elevations (**Fig. 3d**).

Estimating selection on development time, we found no evidence of a significant three-way interaction between development time, species and elevation (*χ^2^*(3)=2.59, P=0.459). However, we did observe a significant development time × elevation interaction (*χ^2^*(3)= 47.33, P<0.001). This suggests that selection on development time changed across elevations, but did not differ between species. Selection favoured faster development for both species at 500m, but slower development at 2,000m (**Table 1**). Selection on development time in the novel environments was therefore in the direction of plasticity for *S. chrysanthemifolius* at 2,000m (slower development at higher elevations), as well as for *S. aethnensis* at 500m (faster development at lower elevations).

**Table 1.**
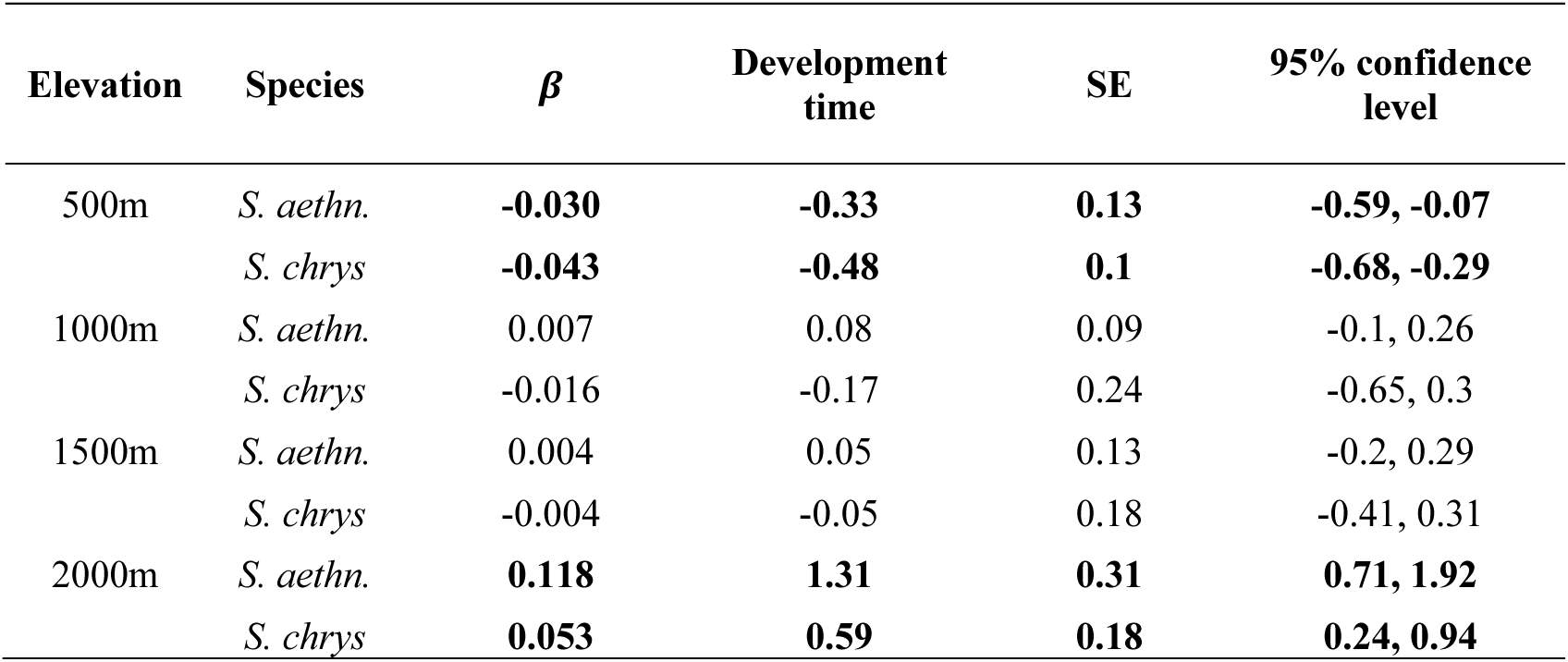
Selection on development time across elevations for both species. Significant estimates of selection are denoted in bold (P<0.05). Selection was only significant at elevational extremes, and in the same direction for both species.

| Elevation | Species | $\beta$ | Development time | SE | 95% confidence level |
| --- | --- | --- | --- | --- | --- |
| 500m | *S. aethn.* | **-0.030** | **-0.33** | **0.13** | **-0.59, -0.07** |
|  | *S. chrys* | **-0.043** | **-0.48** | **0.1** | **-0.68, -0.29** |
| 1000m | *S. aethn.* | 0.007 | 0.08 | 0.09 | -0.1, 0.26 |
|  | *S. chrys* | -0.016 | -0.17 | 0.24 | -0.65, 0.3 |
| 1500m | *S. aethn.* | 0.004 | 0.05 | 0.13 | -0.2, 0.29 |
|  | *S. chrys* | -0.004 | -0.05 | 0.18 | -0.41, 0.31 |
| 2000m | *S. aethn.* | **0.118** | **1.31** | **0.31** | **0.71, 1.92** |
|  | *S. chrys* | **0.053** | **0.59** | **0.18** | **0.24, 0.94** |

### Hypothesis II: G×E across elevation differed for development time and survival

We predicted that stronger patterns of G×E would emerge at elevations further from the home site of each species. We therefore expected genetic variance to increase and genetic correlations between the home site and other elevations to weaken as environments became novel. We observed the predicted pattern for survival. Genetic variance in survival was near zero at the native elevations of both species and significantly greater at the novel environments (**Fig. 4a**). Genetic variance increased consistently with elevation for *S. chrysanthemifolius,* but remained close to zero across most elevations for *S. aethensis* and only increased at 500m (where mean survival was lowest; **Fig. 3c**). For development time, we found the opposite pattern to our original prediction. For both species, genetic variance in development time was more than double at elevations within the native range compared to near-zero genetic variance at the range edge and the novel environment (**Fig. 4b**). Where genetic variance in development time and survival increased significantly, we tested that our observed estimates of genetic variance were statistically significant. We found that our observed estimates exceeded the null distribution for both species and traits (**Fig. S3**), suggesting that the increase in genetic variance (within the native range for development time, and in the novel environment for survival) was not due to random chance, but meaningful differences among genotypes created by G×E.

**Fig. 4.**
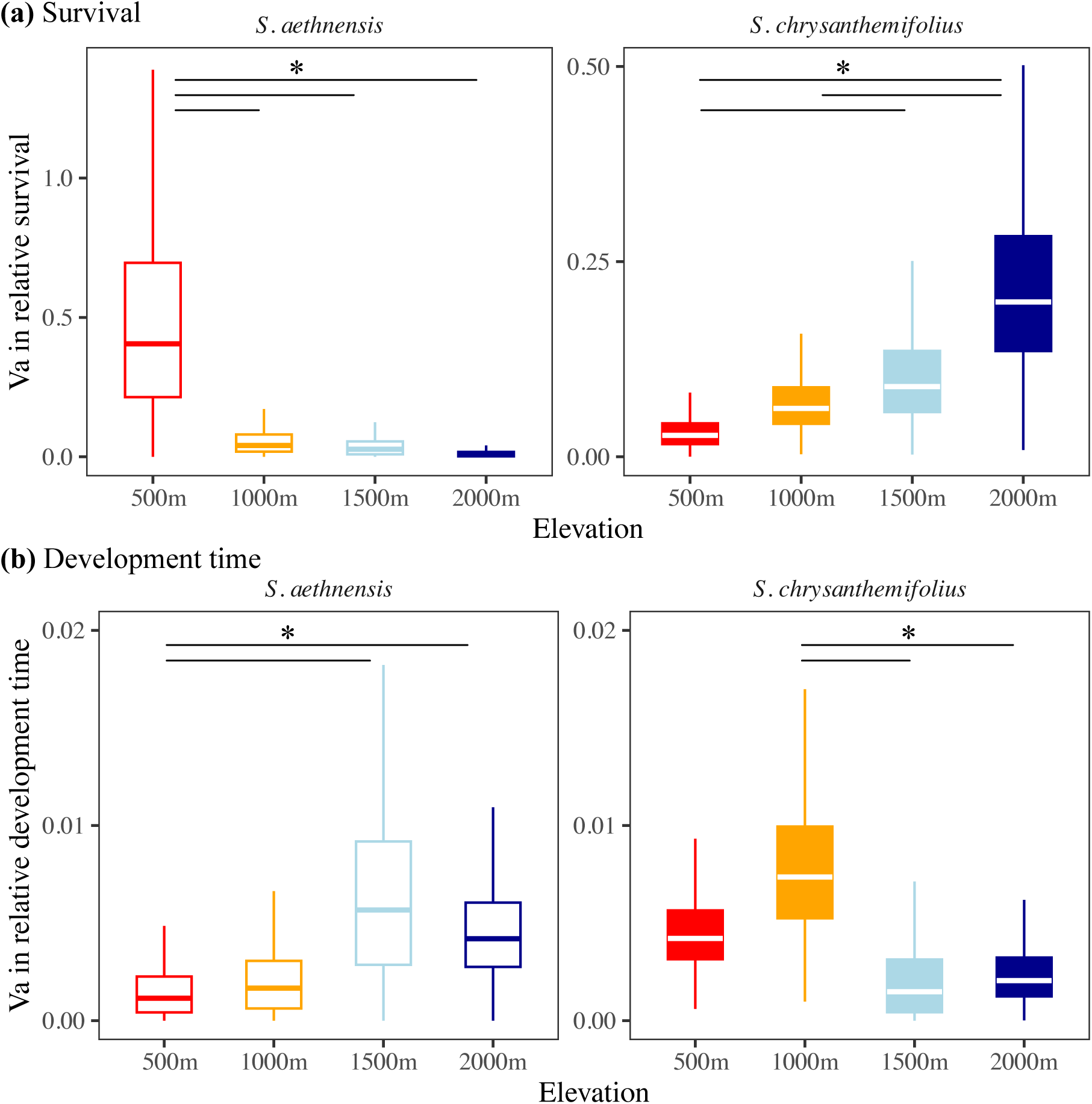
Genetic variance in **(a)** survival and **(b)** development time to seedling establishment for both species at all elevations. **(a)** At their native elevations, genetic variance in survival was near zero, but greater at novel elevations. **(b)** For both species, genetic variance in development time was lower at novel elevations compared to native elevations. Horizontal lines denote elevations that show significant differences in genetic variance whereby <10% of the distributions overlap. >90% of the posterior distributions do not overlap.

Genetic correlations in survival across elevations were all >0.7 for *S. chrysanthemifolius,* and ranged from 0.06-0.58 for *S. aethnensis* (**Table S2**). By contrast, genetic correlations for development time across environments ranged between 0.35-0.79 for both species (**Table S2**). This suggests that rank changes in genotype survival across elevations were likely to have only contributed moderately to G×E in this experiment, and mainly for survival in *S. aethnensis*. However, confidence intervals for the estimates of cross-elevation genetic correlations were often large and overlapping zero, particularly for survival in *S. aethnensis* (**Table S2**), and so care must be taken in their interpretation.

### Hypothesis III: Genetic variation for faster development was associated with higher fitness in novel environments

In our original hypothesis, we predicted that greater G×E in both development time and survival would emerge in novel environments. However, given that development time showed the opposite pattern (a reduction in genetic variance in novel environments), we tested whether genetic variance in development time within the native range was associated with genetic variance in survival at the novel elevations. This was necessary because estimating correlations where genetic variance is near-zero is uninformative. We found that both species showed strong negative genetic correlations between development time within the native range and survival at the novel elevation (**Fig. 5a**). This suggests that genotypes that develop faster in native environments are associated with higher fitness in novel environments (**Fig. 5b**).

**Fig. 5.**
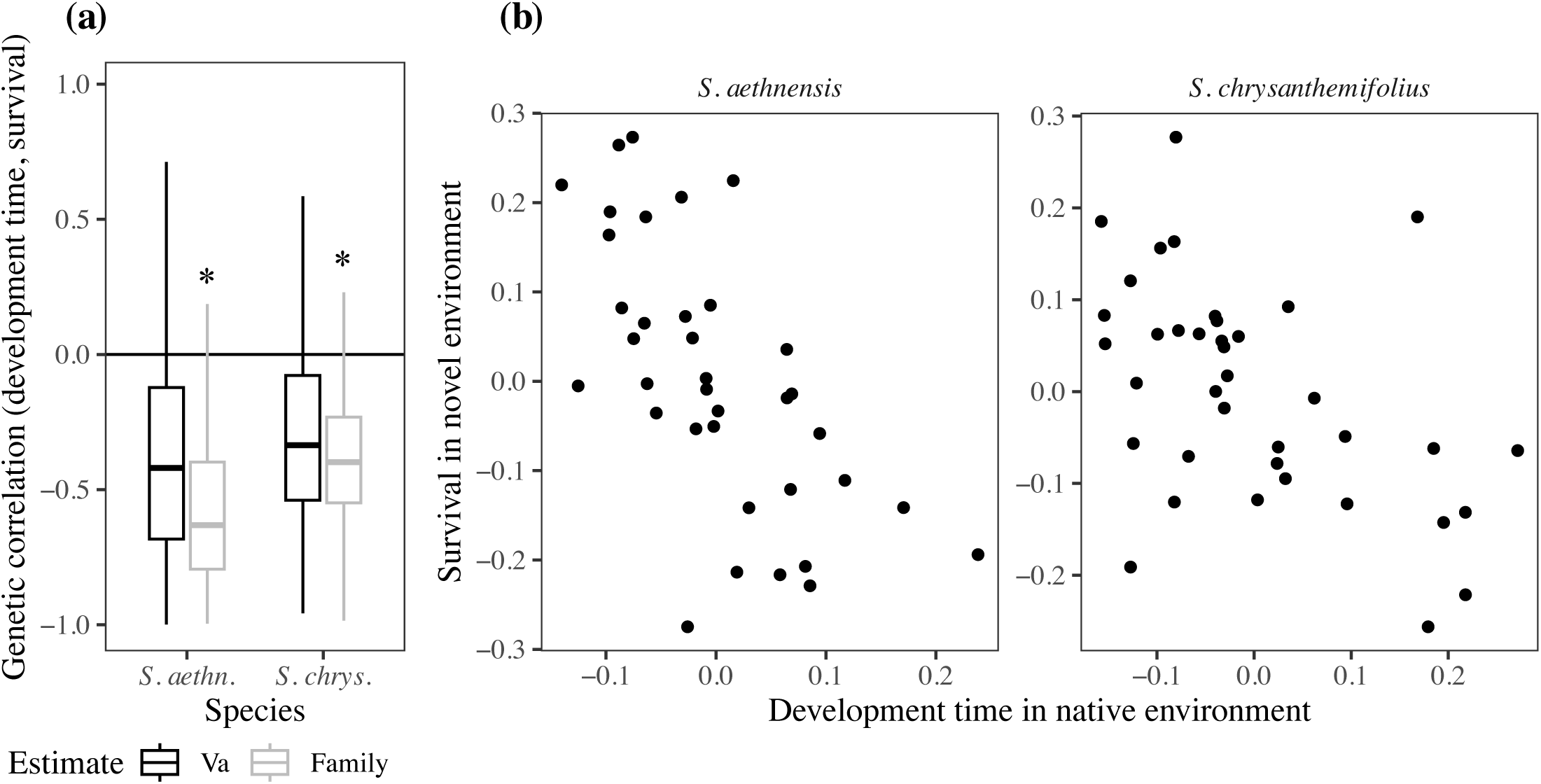
The genetic correlation between development time at a site in the native range and survival in the novel environment. **(a)** Boxplots represent the posterior distribution for the estimates of the genetic correlation, with asterisks denoting correlations where <10% of their distributions overlap. Due to the difficulty of estimating the cross-elevation genetic correlation, we included estimates for the among-family variance, which shows the same pattern but is estimated more precisely due to higher replication of families than sires. **(b)** Mean estimates for the sires (i.e., Best Linear Unbiased Predictors) that represent the genetic correlation across elevations. For both species, genotypes with faster development in the native range showed higher survival in the novel environment.

## Discussion

We reciprocally planted seeds of two ecologically contrasting species of *Senecio* wildflowers across an elevational gradient. By tracking seedling survival and development time to seedling establishment, we found support for three hypotheses that test how adaptive potential of populations change from native to novel environments along an ecological gradient. First, for both species, selection favoured faster development at 500m but slower development at 2000m (**Table 1**). Second, both species showed high survival but low genetic variance in survival in their native elevations, suggesting that all genotypes could maintain similarly high survival within their native range (**Fig. 4a**). However, where mean survival was reduced in novel environments, genetic variance in survival increased, which suggests a greater potential for adaptation to the novel environment (**Fig. 4a**). Surprisingly, genetic variance in development time showed the opposite trend, with greater genetic variance observed within the native range compared to near-zero variance in novel environments (**Fig. 4b**). Third, for both species, we detected negative genetic correlations between development time within the native range and survival in novel environments (**Fig. 5**). Genotypes with faster development times in the native range therefore had greater survival in novel environments, which suggests that genetic variation in early life history traits can predict which genotypes could increase adaptive potential to novel environments. Our results therefore suggest that it could be possible to predict genotypes that could increase the resilience of populations to environmental change, which could benefit conservation strategies, as discussed below.

### Eco-evolutionary inferences of population responses to environmental change

The expression of greater genetic variance in fitness in novel environments was consistent for both *Senecio* species, suggesting that increased adaptive potential in novel environments could be a general pattern for natural populations. Such a pattern is consistent with studies showing that genetic variance in fitness changes depending on the environment (Sheth et al. 2018), increases under drought stress in the glasshouse (Torres-Martínez et al. 2019), and increases in populations suffering population declines (Kulbaba et al. 2019; Walter et al. 2023). However, we still have a poor understanding of the capacity for these genetic differences in relative fitness in novel environments to allow adaptation to environmental change. Further research needs to identify the conditions (e.g., the level of stress and key environmental variables) that lead to increases in genetic variance in fitness. In particular, it would be valuable to quantify the level of stress necessary to induce genetic variance in fitness and then determine whether population persistence is possible under those same stress conditions (i.e., where mean fitness is not so low as to make extinction more likely). Such information can identify the potential not only for adaptation, but also population persistence, to more accurately predict population resilience under rapid environmental change.

### Genetic differences in development costs increase adaptive potential in novel environments

Although stressful environments are expected to release cryptic genetic variation in phenotypes, evidence is rare (McGuigan and Sgrò 2009; Paaby and Rockman 2014). Meta-analyses have found no support for increased genetic variance in novel environments for traits associated with morphology or development (Charmantier and Garant 2005; Noble et al. 2019; Riley et al. 2023). This could be due to inconsistencies as to what studies considered stressful versus novel, and further, the extent of the severity of stressful environments or treatments (Riley et al. 2023). Given that we found opposite patterns for the expression of genetic variance in development time (reduction) versus survival (increase) in novel environments, our results suggest that stress or environmental novelty could have more scope to increase genetic variance for fitness compared to life history traits. Whether these patterns remain if we consider other components of fitness or other novel environments remains to be tested. However, reduced genetic variance for development time in novel environments suggests that adaptive plasticity that helps to maintain fitness within native environments breaks down in novel environments, which could cause variance in fitness to increase. Where plasticity fails to maintain fitness in novel environments, a stress response could emerge that is the same for all genotypes, which increases genetic variance in fitness because genotypes differ in the fitness costs incurred in the novel environment.

Exceeding the limits of plasticity is expected to create fitness costs, but these are notoriously difficult to detect because (1) selection may remove genotypes early in life, and (2) novel environments could produce a canalised stress responses that induces the same phenotype in all individuals (DeWitt et al. 1998; Auld et al. 2010; Murren et al. 2015). In particular, selection is expected to remove genetic variation in life history traits, which could have created low levels of variation in development time (Angert et al. 2014). However, we provide an alternative option – increased genetic variance in fitness in novel environments could be created by genetic variation in the costs of developing in a novel environment (**Fig. 5**). Our results therefore suggest that the genotypes that are better able to develop quickly in their native range suffer less developmental costs, which leads to better survival in novel environments. This is because greater environmental unpredictability can impose a fitness cost to delaying development in a novel environment (Walter et al. 2018), even in environments that favour slower development, such as at 2000m in our study.

### Implications for predicting population and species’ persistence under environmental change

By connecting variation in development within the native range to survival in novel environments, we show that the genetic variation observed within the native range could potentially be used to predict adaptive potential and the fitness responses of genotypes to novel environments. This is encouraging for conservation efforts because it means that it may be possible to identify genotypes that could help populations better cope with environmental change. However, further work needs to quantify how different early life history traits relate to life-time fitness in novel environments, and how consistent this is across taxa and across different environmental stresses. Future research should target key traits and life history stages within native environments, and test whether they could be used to identify genotypes that will be more resilient to environmental change (Bay et al. 2017). Using genomic prediction could then be used to identify adaptive genetic variation for novel environments created by climate change (Waldvogel et al. 2020; Ashraf et al. 2022). Such an approach can increase our ability to predict the adaptive capacity of populations facing environmental change as well as enhance conservation efforts seeking to identify genotypes that could aid population persistence. Furthermore, incorporating estimates of genetic diversity can test how demographic history determines the expression of genetic variance in novel environments, and in particular, identify whether populations with low genetic diversity simply lack the genotypes necessary to aid population persistence in novel environments, and whether such genotypes can then be acquired from elsewhere in the geographical range (Sgrò et al. 2011; Olazcuaga et al. 2023). Such information will be crucial for developing techniques for genetic rescue of threatened populations or species.

### Sources of G×E in development time and survival

Our results suggest that the emergence of cryptic genetic variation underlies increased adaptive potential in novel environments that we observed (McGuigan and Sgrò 2009; Paaby and Rockman 2014). However, we still do not know how epigenetic or gene regulation differences among genotypes underlie increased adaptive potential in novel environments (McGuigan et al. 2021; Liu et al. 2022). It is possible that increased genetic variance in fitness could be linked to conditionally neutral alleles that have zero fitness effects in native environments, but strong fitness effects in a novel environment because they have not been exposed to selection that would have otherwise removed them (Paaby and Rockman 2014). We therefore need further experiments that identify the genetic and environmental mechanisms that increase adaptive potential in stressful or novel environments.

### Limitations of inferring adaptive potential using seed experiments

We provide strong evidence in two *Senecio* species that moving into novel environments where plasticity can no longer maintain fitness, genetic variation in fitness increases. This is consistent with our previous study that showed greater genetic variance in fitness measured as reproductive (flower) investment for *S. chrysanthemifolius* at the novel 2000m elevation. However, although tracking seedlings allows key aspects of early life traits to be included in our current study, we were unable to measure performance at later life history stages to capture traits related to lifetime fitness. This requires tracking plants from seeds to maturity and fecundity (e.g., Shaw et al. 2008; Sheth et al. 2018; Kulbaba et al. 2019; Peschel et al. 2020), which is a challenge for field experiments, but vital for understanding the potential for adaptation at range margins and in response to environmental change (Bridle and Hoffmann 2022). Our genetic correlations between development time and survival were not significant for sire variance (but were for among-family variance), suggesting that further replication is needed to more precisely estimate genetic correlations across environments, particularly for different traits. Our estimates of genetic variance in development time also exclude seedlings that died early or failed to germinate, and so selection could have removed genetic variation that could have led to greater genetic variance in development time in novel environments, as originally predicted. However, this should only be an issue for *S. chrysanthemifolius* because selection on *S. aethnensis* at low elevations largely occurred after seedling establishment, meaning that development time was measured for most of the emergent seedlings. Future experiments should focus on estimating developmental and morphological traits in seedlings well before establishment (i.e., before selection removes variation) to better quantify how plasticity and adaptive potential could contribute to population persistence in novel environments.

### Conclusions

Using an extensive reciprocal planting of two contrasting *Senecio* wildflower species across an elevation gradient, we show that increased genetic variance in fitness in novel environments increases adaptive potential. We also show that G×E in early development showed the opposite trend, with greater genetic variance in seedling development time observed in native compared to novel environments. A strong negative genetic correlation between development time within the native range with fitness outside the range suggests that genotypes that develop faster within the range show greater fitness in novel environments. While further work needs to understand the mechanisms underlying this pattern, these data suggest that genetic variance in life history traits could predict genotypes that could aid population persistence in novel environments. This information should be useful for efforts to predict population and community persistence under rapid environmental change, and to identify genotypes that could help persistence and adaptation to novel environments created by global change.

## Acknowledgements

We are very grateful to Piante Faro (Giarre, Italy) for providing us with glasshouse facilities. We thank Giuseppe Riggio for generously providing us access to the 1,000m field site. This work was carried out using the computational facilities of the Advanced Computing Research Centre, University of Bristol.

## Author contributions

GW designed the study with AC, SH, SC and JB. GW, DT, ES, MM, GP and AC conducted the experiments, collected and curated the data. GW analysed the data and wrote the paper with help from KM, AW, JC, SP, and input from all authors.

## Data availability

Data is available at https://doi.org/10.5061/dryad.k6djh9wdf.

