## Supplementary material for "Increased adaptive potential in novel environments can be predicted from genetic variance in development time expressed in native environments"

**Contents**

**Table S1** Sampling locations
**Fig. S1** Map of sampling locations **Fig. S2** Demography during the field experiment
**Fig. S3** Testing genetic variance
**Table S2** Genetic (co)variance matrices

**Table S1:** Location of sampled individuals for the parental generation of the breeding design for each species. The final two column denote the number of individuals used as sires and dams in the breeding design.

| **Species** | **Site** | **Elevation** | **Latitude** | **Longitude** | **# Sires** | **# Dams** |
| --- | --- | --- | --- | --- | --- | --- |
| *S. aethn.* | Etna South | 2,600m | 37°43'13.28"N | 15° 0'3.54"E | 1 | 1 |
|  |  | 2,500m | 37°43'3.80"N | 14°59'59.20"E | 8 | 7 |
|  |  | 2,400m | 37°42'46.50"N | 14°59'41.30"E | 5 | 5 |
|  |  | 2,200m | 37°42'24.82"N | 14°59'42.69"E | 2 | 5 |
|  | Etna North | 2,600m | 37°46'39.90"N | 15° 0'23.00"E | 8 | 4 |
|  |  | 2,500m | 37°46'53.70"N | 15° 0'28.80"E | 3 | 2 |
|  |  | 2,400m | 37°47'7.46"N | 15° 0'35.05"E | 4 | 8 |
|  |  | 2,200m | 37°47'32.82"N | 15° 1'14.53"E | 5 | 3 |
|  | **Totals** |  |  |  | **36** | **35** |
| *S. chrys.* | Bonnano | 790m | 37°38'24.92"N | 15° 2'50.80"E | 9 | 11 |
|  | Cacciola | 680m | 37°37'31.32"N | 15° 3'26.71"E | 7 | 6 |
|  | Poggofelice | 590m | 37°39'44.31"N | 15° 5'48.55"E | 6 | 6 |
|  | Spina | 730m | 37°39'19.27"N | 15° 4'30.92"E | 10 | 10 |
|  | Trecastagni | 570m | 37°36'46.67"N | 15° 4'29.64"E | 6 | 5 |
|  | **Totals** |  |  |  | **38** | **38** |

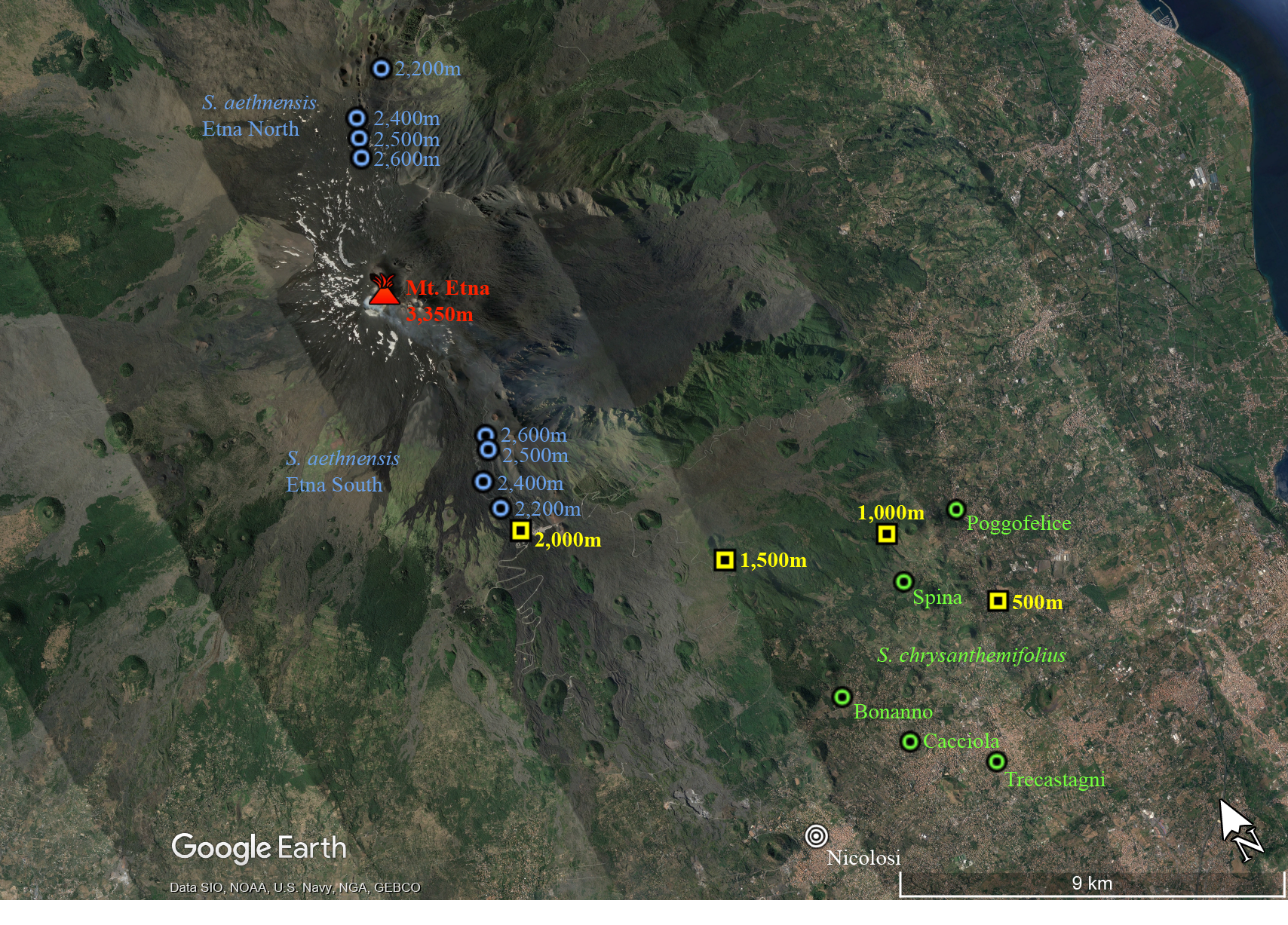

**Fig. S1** Map with the locations of transplant sites (yellow), and the sites that the parental genotypes were sampled from for both *S. aethnensis* (blue) and *S. chrysanthemifolius* (green).

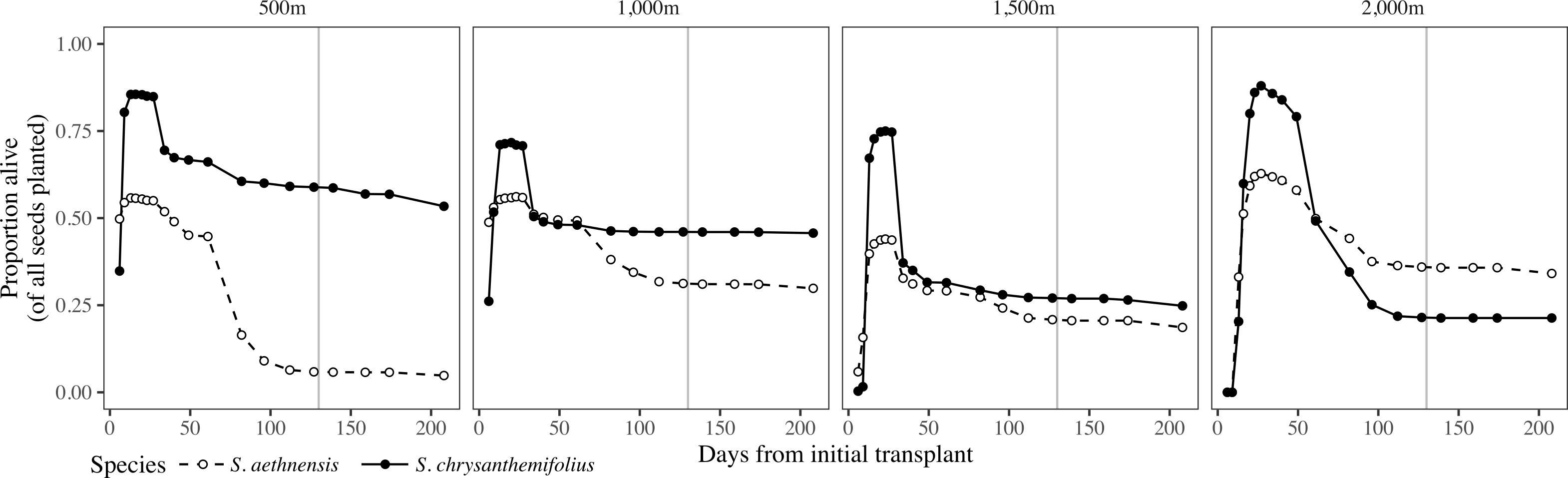

**Fig. S2** Proportion of plants alive at each census day, for each transplant elevation. Filled circles and solid lines represent *S. chrysanthemifolius*, while unfilled circles and broken lines represent *S. aethnensis*. Vertical grey lines represent when mortality stabilised at the end of summer.

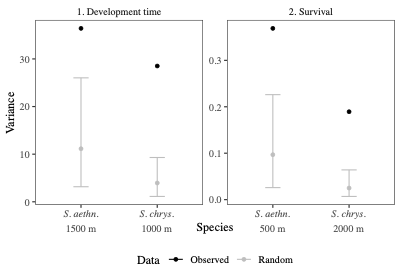

**Fig. S3** Observed estimates of genetic variance (black circles) are greater than the null distribution (grey circles and 95% HPD interval) for both traits and species, suggesting that the increase in genetic variance highlighted in the main text (and in **Fig. 4**) capture statistically significant genetic variance. To construct the null distributions, we estimated genetic variance using data that was randomized with respect to family identification. To maintain differences among experimental blocks in the field, we randomized offspring within each block separately. At each elevation, we conducted 200 randomizations of the data, reapplied equation 3, and saved the mean estimate of genetic variance for each model. Where the mean of the observed model (in black) exceeds the null distribution (in grey), there is evidence that our estimate of genetic variance is greater than expected under random sampling, and is therefore statistically significant.

**Table S2** Genetic variances at each elevation (along diagonal in grey) and genetic correlations across elevations (lower diagonal) for **(a)** survival, and **(b)** development time. Numbers in parentheses denote the 90% Highest Posterior Density (HPD) interval for each parameter estimated.

|  | **(a)** Survival | | | | | | | | | | |
| --- | --- | --- | --- | --- | --- | --- | --- | --- | --- | --- | --- |
|  | | *S. aethnensis* | | | |  |  | *S. chrysanthemifolius* | | | |
|  | | 500m | 1000m | 1500m | 2000m |  |  | 500m | 1000m | 1500m | 2000m |
| 500m | | 0.519  (0, 1.022) |  |  |  |  | 500m | 0.033 (0.004, 0.064) |  |  |  |
| 1000m | | 0.58  (0.19, 0.95) | 0.059  (0, 0.134) |  |  |  | 1000m | 0.74  (0.48, 0.98) | 0.071 (0.013, 0.122) |  |  |
| 1500m | | 0.39  (-0.14, 0.96) | 0.4  (-0.17, 0.96) | 0.04  (0, 0.096) |  |  | 1500m | 0.76  (0.53, 0.98) | 0.84  (0.67, 0.99) | 0.102 (0.014, 0.191) |  |
| 2000m | | 0.08  (-0.59, 0.89) | 0.06  (-0.68, 0.83) | 0.12  (-0.64, 0.82) | 0.014  (0, 0.036) |  | 2000m | 0.75  (0.51, 0.98) | 0.83 (0.66, 1) | 0.82  (0.63, 0.99) | 0.222 (0.044, 0.393) |
|  | **(b)** Development time | | | | | | | | | | |
|  | | *S. aethnensis* | | | |  |  | *S. chrysanthemifolius* | | | |
|  | | 500m | 1000m | 1500m | 2000m |  |  | 500m | 1000m | 1500m | 2000m |
| 500m | | 0.002  (0, 0.004) |  |  |  |  | 500m | 0.005  (0.002, 0.008) |  |  |  |
| 1000m | | 0.35  (-0.29, 0.97) | 0.002  (0, 0.005) |  |  |  | 1000m | 0.79  (0.59, 0.98) | 0.008  (0.002, 0.013) |  |  |
| 1500m | | 0.39  (-0.23, 0.94) | 0.42  (-0.14, 0.96) | 0.007  (0, 0.013) |  |  | 1500m | 0.43  (-0.16, 0.99) | 0.42  (-0.26, 0.97) | 0.002  (0, 0.005) |  |
| 2000m | | 0.46  (-0.11, 0.98) | 0.54 (0.02, 0.97) | 0.48 (0.01, 0.96) | 0.005 (0.001, 0.008) |  | 2000m | 0.72  (0.44, 0.99) | 0.65  (0.29, 0.99) | 0.39  (-0.23, 0.96) | 0.002  (0, 0.005) |
